# Diploid chromosome-scale assembly of the *Muscadinia rotundifolia* genome supports chromosome fusion and disease resistance gene expansion during *Vitis* and *Muscadinia* divergence

**DOI:** 10.1101/2020.06.02.119792

**Authors:** Noé Cochetel, Andrea Minio, Mélanie Massonnet, Amanda M. Vondras, Rosa Figueroa-Balderas, Dario Cantu

## Abstract

*Muscadinia rotundifolia*, the muscadine grape, has been cultivated for centuries in the southeastern United States. *M. rotundifolia* is resistant to many of the pathogens that detrimentally affect *Vitis vinifera*, the grape species commonly used for winemaking. For this reason, *M. rotundifolia* is a valuable genetic resource for breeding. Single-molecule real-time reads were combined with optical maps to reconstruct the two haplotypes of each of the 20 *M. rotundifolia* cv. Trayshed chromosomes. The completeness and accuracy of the assembly were confirmed using a high-density linkage map of *M. rotundifolia.* Protein-coding genes were annotated using an integrated and comprehensive approach. This included using Full-length cDNA sequencing (Iso-Seq) to improve gene structure and hypothetical spliced variant predictions. Our data strongly support that *Muscadinia* chromosomes 7 and 20 are fused in *Vitis* and pinpoint the location of the fusion in Cabernet Sauvignon and PN40024 chromosome 7. Disease-related gene numbers in Trayshed and Cabernet Sauvignon were similar, but their clustering locations were different. A dramatic expansion of the Toll/Interleukin-1 Receptor-like Nucleotide-Binding Site Leucine-Rich Repeat (TIR-NBS-LRR) class was detected on Trayshed chromosome 12 at the *Resistance to Uncinula necator 1* (*RUN1*)/ *Resistance to Plasmopara viticola 1* (*RPV1*) locus, which confers strong dominant resistance to powdery and downy mildews. A genome browser for Trayshed, its annotation, and an associated Blast tool are available at www.grapegenomics.com.

## INTRODUCTION

Viticulture and winemaking typically make use of *Vitis vinifera* fruit. Though *V. vinifera* wines have desirable organoleptic properties, *V. vinifera* is extremely sensitive to diverse pathogens. In contrast, *Muscadinia* is a genus closely related to *Vitis* (Small 1913; Bouquet 1978; Olmo 1986; Olien 1990) that is resistant to diverse biotic and abiotic stresses. *Muscadinia rotundifolia* is the foundation of the muscadine grape industry in the United States of America (USA), where its fruit, juice, and wine are produced (Olien 1990). *M. rotundifolia* is native to the warm and humid southeastern USA. Its natural habitat includes the states bordering the Gulf of Mexico and extends from northern Arkansas to Delaware (Bouquet 1978; Olmo 1986; Olien 1990; Heinitz *et al.* 2019).

*Vitis* and *Muscadinia* belong to Vitaceae, a family that contains 16 other genera and approximately 950 species (Wen *et al.* 2018). These genera are estimated to have diverged between 18 and 47 million years ago (Ma) (Wan *et al.* 2013; Liu *et al.* 2016; Ma *et al.* 2018), a process that involved several chromosome rearrangements (Karkamkar *et al.* 2010; Wen *et al.* 2018). *Vitis* has 19 chromosomes (2n = 38) (Olmo 1937), but like other genera in Vitaceae, including *Ampelopsis, Parthenocissus*, and *Ampelocissus, Muscadinia* has 20 chromosomes (2n = 40) (Branas 1932; Patil and Patil 1992; Karkamkar *et al.* 2010; Chu *et al.* 2018). Though successful crosses yielding fertile hybrids have been obtained, early attempts at producing hybrids of *Vitis* and *Muscadinia* often resulted in sterile progeny; this and the graft incompatibility observed between the two genera (Patel and Olmo 1955) is thought to be caused by their difference in chromosome number (Ravaz 1902; Dearing 1917; Patel and Olmo 1955; Olmo 1971; Bouquet 1980; Walker *et al.* 1991). SSR markers and genetic maps revealed that linkage groups LG7 and LG20 in *Muscadinia* correspond to the proximal and distal regions of chromosome 7 in *Vitis* (Blanc *et al.* 2012; Delame *et al.* 2019; Lewter *et al.* 2019). Chromosome 7 in *Vitis* may be derived from the fusion of its ancestor’s chromosomes 7 and 20. The presence of residual telomeric repeats in chromosome 7 of *Vitis* (Lewter *et al.* 2019) could provide additional support for this hypothesis, insight into the evolutionary history of Vitaceae, and understanding of the structure and function of the *Muscadinia* genome.

*M. rotundifolia* is a desirable partner with which to hybridize because it is resistant to many stresses, pests, and diseases that adversely affect *V. vinifera* and cause substantial crop loss (Olmo 1971). *M. rotundifolia* is resistant to Pierce’s disease *(Xylella fastidiosa)* (Ruel and Walker 2006), phylloxera *(Daktulosphaira vitifolia)* (Ravaz 1902; Davidis and Olmo 1964; Firoozabady and Olmo 1982), downy mildew (*Plasmopara viticola*) (Olmo 1971; Staudt and Kassemeyer 1995), powdery mildew *(Erysiphe necator* syn. *Uncinula necator)* (Olmo 1986; Merdinoglu *et al.* 2018), and other diseases and pests (Ravaz 1902; Olmo 1971; Walker *et al.* 2014). Several loci associated with resistance to pathogens affecting *V. vinifera* were identified in *M. rotundifolia*, including *Resistance to Uncinula necator 1 (RUN1)* (Pauquet *et al.* 2001), *RUN2* (Riaz *et al.* 2011), *Resistance to Erysiphe Necator 5 (REN5)* (Blanc *et al.* 2012), *Resistance to Plasmopara viticola 1 (RPV1)* (Merdinoglu *et al.* 2003), and *RPV2* (Merdinoglu *et al.* 2018). However, the lack of a *M. rotundifolia* reference sequence and gene annotation limits gene discovery and the characterization of resistance loci.

A draft assembly of the *M. rotundifolia* cv. Trayshed (Trayshed, hereafter) genome using Single-Molecule Real-Time (SMRT) sequencing was instrumental in resolving the genetic basis of sex determination in grapes (Massonnet *et al.*, 2020). Here, we report the phased, chromosome-scale assembly of Trayshed, which was produced improving the previous SMRT assembly with the introduction of the optical maps in a hybrid scaffolding approach and consensus genetic map from multiple wild and cultivated grape species. Full-length cDNA isoforms (Iso-Seq) were also sequenced and used as transcriptional evidence for the annotation of protein-coding genes. This assembly and its annotation were used to identify where *M. rotundifolia* chromosomes 7 and 20 fused to create *V. vinifera* chromosome 7 but also to identify genes at the *RUN1*/*RPV1* locus.

## MATERIALS AND METHODS

### Trayshed chromosome construction

A Saphyr Genome Imaging Instrument was used to generate a Bionano Next-Generation Mapping (NGM) of Trayshed. Ultra-high molecular weight DNA (> 500 kbp) was extracted from young leaves by Amplicon Express (Pullman, WA). DNA was then labeled with a DLE-1 non-nicking enzyme (CTTAAG) and stained according to the Bionano Prep™ Direct Label and Stain (DLS) Kit (Bionano Genomics, San Diego, CA) instructions. Labeled DNA was loaded onto the SaphyrChip nanochannel array for imaging on the Saphyr system (Bionano Genomics, San Diego, CA). Imaged molecules longer than 150 kbp were kept. These molecules were then assembled using Bionano Solve v.3.3 (Lam *et al.* 2012) with the following parameters “haplotype”, “noES”, and “noCut”, significance cutoffs were adopted *P* < 1e-10 to generate draft consensus contigs, *P* < 1e-11 for draft consensus contig extension, *P* < 1e-15 for the final merging of the draft consensus contigs. The complete configuration file used for the assembly is available as **Supplemental File S1**. This assembly procedure produced a 1.18 Gbp consensus genome map with an N50 of 5.6 Mbp.

PacBio contigs generated with FALCON-Unzip as described in Massonnet *et al*. (2020) were scaffolded with the genome maps using HybridScaffold v.04122018 (Lam *et al.* 2012) with options “-B2 -N1” for conflict resolution. The complete parameter configuration is available as **Supplemental File S2**. The ploidy state of the 2,000 rhAmpseq markers conserved across wild and domesticated grapes (Zou *et al.* 2020) was evaluated using DISPR (https://github.com/douglasgscofield/dispr) to keep the ones found in a maximum of two copies across different sequences. Hybrid scaffolds were then pre-processed to prevent marker duplication and wrong primary contigs/haplotigs co-placement. Two sets of chromosome-scale pseudomolecules were reconstructed using the pre-processed hybrid scaffolds and HaploSync v.1 tool suite (https://github.com/andreaminio/HaploSync). The hybrid scaffolds were sorted using the curated rhAmpseq markers (Zou *et al.* 2020). Sequences were treated independently from FALCON Unzip’s classification as primary contigs or haplotigs but, the relationship information between the primary contigs and the corresponding alternate haplotigs was conserved during the process. Then, the two haplotypes of each chromosome were compared to identify and correct assembly errors and fill gaps.

### cDNA library preparation and sequencing

Total RNA from Trayshed leaves were isolated using a Cetyltrimethyl Ammonium Bromide (CTAB)-based extraction protocol as described in Blanco-Ulate *et al.* (2013). RNA purity was evaluated with a Nanodrop 2000 spectrophotometer (Thermo Scientific, Hanover Park, IL), RNA quantity with a Qubit 2.0 Fluorometer and a broad range RNA kit (Life Technologies, Carlsbad, CA), and RNA integrity by electrophoresis and an Agilent 2100 Bioanalyzer (Agilent Technologies, CA). Total RNA (300 ng, RNA Integrity Number > 8.0) were used for cDNA synthesis and library construction. A cDNA SMRTbell library was prepared. First-strand synthesis and cDNA amplification were accomplished using the NEB Next Single Cell/Low Input cDNA Synthesis & Amplification Module (New England, Ipswich, MA, US). The cDNAs were subsequently purified with ProNex magnetic beads (Promega, WI) following the instructions in the Iso-Seq Express Template Preparation for Sequel and Sequel II Systems protocol (Pacific Biosciences, Menlo Park, CA). ProNex magnetic beads (86 μL) were used to select amplified cDNA >= 2 kbp. At least 80 ng of the size-selected, amplified cDNA were used to prepare the cDNA SMRTbell library. This was followed by DNA damage repair and SMRTbell ligation using the SMRTbell Express Template Prep Kit 2.0 (Pacific Biosciences, Menlo Park, CA) following the manufacturer’s protocol. One SMRT cell was sequenced on the PacBio Sequel I platform (DNA Technology Core Facility, University of California, Davis). IsoSeq reads were extracted using IsoSeq v3 protocol with default parameters and low-quality isoform polished wit LSC v.2.0 (Au *et al.*, 2012). A cDNA library was also prepared using the Illumina TruSeq RNA sample preparation kit v.2 (Illumina, CA, USA) following Illumina’s low-throughput protocol. This library was evaluated for quantity and quality with the High Sensitivity chip in an Agilent 2100 Bioanalyzer (Agilent Technologies, CA) and sequenced in 100 bp paired-end reads using an Illumina HiSeq4000 sequencer (DNA Technology Core Facility, University of California, Davis).

### Genome annotation

The completeness of the gene space in the hybrid assembly was estimated using BUSCO v.3.0.2 (Waterhouse *et al.*, 2018) with the embryophyta_odb9 conserved gene set and by aligning the PN40024 V1 CDS against the Trayshed assembly using BLAT v.36×2 with default parameters (Kent 2002). PN40024 CDS were filtered before the alignment and included only single-copy genes (*i.e*., with unique mapping on its genome). The structural annotation of the *M. rotundifolia* cv. Trayshed genome was performed with a modified version of the pipeline used for the *Vitis vinifera* cv. Zinfandel genome (Vondras *et al.* 2019) and fully described here: https://github.com/andreaminio/AnnotationPipeline-EVM_based-DClab. Briefly, high-quality Iso-Seq Trayshed data were used to produce high-quality gene models for training gene predictors in PASA v.2.3.3 (Haas 2003) along with transcript evidence obtained from RNAseq data by performing transcriptome assemblies using Stringtie v.1.3.4d (Pertea *et al.* 2015) and Trinity v.2.6.5 (Grabherr *et al.* 2011) and from external databases. Public databases, transcriptome assemblies, and the Iso-Seq data described above were used as transcript and protein evidence. They were mapped on the genome using PASA v.2.3.3 (Haas 2003), MagicBLAST v.1.4.0 (Boratyn *et al.* 2019), and Exonerate v.2.2.0 (Slater and Birney 2005). *Ab initio* predictions were generated using BUSCO v.3.0.2 (Waterhouse *et al.*, 2018), Augustus v.3.0.3 (Stanke *et al.* 2006), GeneMark v.3.47 (Lomsadze 2005), and SNAP v.2006-07-28 (Korf 2004). Repeats were annotated using RepeatMasker v.open-4.0.6 (Smit *et al.* 2013). Next, EvidenceModeler v.1.1.1 (Haas *et al.* 2008) used these predictions to generate consensus gene models. The final functional annotation was produced combining blastp v.2.2.28 (Camacho *et al.* 2009) hits against the Refseq plant protein database (ftp://ftp.ncbi.nlm.nih.gov/refseq, retrieved January 17th, 2019) and InterProScan v.5.28-67.0 (Jones *et al.* 2014) outputs using Blast2GO v.4.1.9 (Gotz *et al.* 2008). To evaluate hemizygosity, CDS from each haplotype were aligned on the opposite haplotype using GMAP v. 2019-09-12 (Wu and Watanabe 2005) and alignments with at least 80% identity and coverage were considered valid. To prevent over-estimation of the gene content diversity between the two haplotypes due to pseudomolecule reconstruction partiality, unplaced sequences were also searched for copies of the CDS missing in the alternative haplotype.

### Genome size quantification by flow cytometry

DNA content was estimated using flow cytometry (n = 3 individual leaves). *Lycopersicon esculentum* cv. Stupické polní tyčkové rané DNA was selected as the internal reference standard with a genome size of 2C = 1.96 pg; 1C = 958 Mbp (Doležel *et al.* 1992). Nuclei extraction was performed using the Cystain PI absolute P kit (Sysmex America Inc). Approximately 5 mg (0.7 cm^2^) of young healthy leaves from grapevine and tomato were finely cut with a razor blade in a Petri dish containing 500 μL of extraction buffer. The nuclei suspension was filtered through a 50 μm filter (CellTrics, Sysmex America Inc) and 2 mL of a propidium iodide staining solution was added (Doležel and Bartoš 2005; Bertier *et al.* 2013). Measurements were acquired using a Becton Dickinson FACScan (Franklin Lakes, New Jersey) equipped with a 488 nm laser. The data were analyzed using FlowJo v.10 (https://www.flowjo.com/solutions/flowjo). DNA content was inferred by linear regression using the tomato DNA reference standard.

### Comparison of genome assembly and synteny analysis

The Trayshed and Cabernet Sauvignon (Massonnet *et al.*, 2020) genomes were compared with NUCmer from the MUMmer v.4.0.0beta5 tool suite (Marçais *et al.* 2018), and the “--mum” parameter set. Descriptive statistics of the alignment were obtained using the MUMmer script dnadiff. For visualization purposes, alignments with at least 90% identity and 7500 bp long were kept. Blastp v.2.2.28 (Camacho *et al.* 2009) was used to align the annotated proteins of Trayshed and Cabernet Sauvignon. Alignments with at least 80% reciprocal identity and coverage were retained. Pairwise protein information was associated with genes and processed with McScanX v.11.Nov.2013 (Wang *et al.* 2012) with default parameters to identify syntenic regions.

### Localization of *Muscadinia* SNP markers

To localize the Trayshed genomic sequences the corresponding position of each marker identified from *Muscadinia* populations in Lewter *et al* 2019, the sequence surrounding each of them was extracted from PN40024 V0 (Lewter *et al.*, 2019) keeping five hundred base-pair of overhang upstream and downstream of the SNP position. Sequences were then aligned with BLAT v.36×2 (Kent 2002) over the Trayshed chromosomes to recover the marker position. Alignments with at least 90% identity for both query and reference, 90% of coverage on the query, and a maximum of 50-bp-long gaps were considered as hits. They were then filtered for uniqueness based on the number of bases matching. The previously published consensus map of *M. rotundifolia* was used to assess the completeness of the chromosome reconstruction (Lewter *et al.* 2019). Sliding windows (window size = 20 markers, sliding = 10 markers) were designed to move across the uniquely mapped markers on the genome. The percentage of relative marker positions consistent with their genetic position on the high-density genetic map was calculated per window.

### Telomeric repeats analysis

Telomeric repeats of “CCCTAA” and its reverse complement were searched along chromosome 7 of Cabernet Sauvignon and PN40024 V2 (Canaguier *et al.* 2017) using vmatchPattern from the R package Biostrings v.2.56.0 (Pagès *et al.* 2019). After peaks in the distribution of telomeric repeats were identified in both genomes, genomic regions of Cabernet Sauvignon (chr7: 18,675,000-18,677,000 bp) and PN40024 (chr7: 17,572,000 - 17,574,000bp) were extracted. Motif enrichment analysis was performed using MEME v.5.1.0 (Bailey and Elkan 1994) and reported as “logo” using the ggseqlogo R package v.0.1 (Wagih 2017).

### Identification of NBS-LRR genes

Predicted Trayshed and Cabernet Sauvignon proteins were scanned using hmmsearch from HMMER v.3.3 (http://hmmer.org/) with a significance threshold (sequence E-value) of 0.001 and Hidden Markov Models (HMM) corresponding to the following different Pfam (El-Gebali *et al.* 2019) domains: NB-ARC (Pfam PF00931), TIR (PF01582) and LRR (PF00560, PF07723, PF07725, and PF12799). The proteome was also scanned with NLR-annotator (Steuernagel *et al.*, 2020). Coiled-coil (CC) domain-containing proteins were identified by COILS (Lupas *et al.* 1991) during the InterProScan annotation. The identified proteins were then divided into six protein classes according to their domain composition: CC-NBS-LRR, CC-NBS, TIR-NBS-LRR, TIR-NBS, NBS-LRR, and NBS. To capture the largest number of potential NBS-LRR related genes, genes lacking the NB-ARC domain but with “NBS-LRR” functional annotations were also selected and divided into two classes: TIR-X and CC-X. For these eight protein classes, Multiple EM for Motif Elicitation (MEME) analysis was performed with the flags “-mod anr -nmotifs 20” to identify conserved domains for each class. Then, FIMO v.5.1.0 (Find Individual Motif Occurrences) (Grant *et al.* 2011) was run on each protein class using the corresponding MEME results; proteins with at least five conserved domains were kept. NBS-LRR gene clusters were defined as groups of at least two NBS-LRR genes, each separated by no more than eight non-NBS-LRR genes (Richly *et al.* 2002) in a region spanning a maximum of 200 kbp (Holub 2001).

### *RUN1/RPV1* locus analysis and NBS-LRR genes phylogeny

Boundaries of the *RUN1/RPV1* locus were defined by mapping two SSR markers, VMC4f3.1 and VMC8g9 (Barker *et al.* 2005), on Trayshed and Cabernet Sauvignon haplotypes using GMAP v. 2019-09-12 (Wu and Watanabe 2005). Trayshed chromosome 12 was split into 50-kbp blocks and queried with the genomic region corresponding to the locus in Cabernet Sauvignon using tblastx (BLAST v.2.2.29) (Camacho *et al.* 2009). Four-way pairwise comparisons between haplotypes were performed, results were filtered keeping alignments with at least 90% of identity, at least 100 bp long, and located within the boundaries of the resistance locus in the query and the reference. Hits overlapping annotated LTR retrotransposons were discarded. Proteins corresponding to the first alternative spliced variant of each NBS-LRR gene at the *RUN1/RPV1* locus were aligned using MUSCLE v.3.8.31 (Edgar 2004) with default parameters and by including *GAPDH* (*VIT 17s0000g10430*) ortholog sequences as an outgroup. Distances between the alignments were extracted using the R package seqinr v.4.2.4 (Charif and Lobry 2007). The estimation of the corresponding neighbor-joining tree, its rooting, and bootstrapping (1000 replicates) were performed using the R package ape v.5.4.1 (Paradis and Schliep 2019). The tree was drawn using ggtree v.2.2.4 (Yu *et al.* 2017). Species-specific clusters were identified as five or more grouped proteins from the same NBS-LRR class resulting from the phylogenetic analysis with a maximum of one protein coming from the other species in the same tree branch. Coding sequences of the Trayshed-specific protein cluster were aligned per class using MACSE v.2.03 (Ranwez *et al.* 2018) and trimmed using the option “min_percent_NT_at_ends=0.80”. Synonymous substitution rates (dS) were obtained using yn00 from PAML v.4.9f (Yang 2007).

### Data analysis and visualization

Data were parsed with R v.4.0.3 (R Core Team 2020) in RStudio v.1.4.869 (RStudio Team 2020) using tidyverse v.1.3.0 (Wickham *et al.* 2019), GenomicFeatures v.1.40.1 for genomic ranges manipulation (Lawrence *et al.* 2013), and rtracklayer for GFF files v.1.48.0 (Lawrence *et al.* 2009).

### Data availability

Sequencing data are accessible at the NCBI repository under the accessions PRJNA635946 and PRJNA593045. Raw optical maps are available at Zenodo under the DOI 10.5281/zenodo.3866087. The supplemental files are available at Figshare. The pipeline for the gene annotation is available at https://github.com/andreaminio/AnnotationPipeline-EVM_based-DClab. Assembly and annotation files are available at www.grapegenomics.com, which also hosts a genome browser and a blast tool for Trayshed.

## RESULTS AND DISCUSSION

### Construction of the twenty phased *Muscadinia rotundifolia* cv. Trayshed chromosomes

The haploid genome size of *M. rotundifolia* cv. Trayshed evaluated by flow cytometry was estimated at 483.4 ± 3.1 Mbp, a size comparable to PN40024 (487 Mbp; Jaillon et al. 2007), but smaller than Cabernet Sauvignon (557.0 ± 2.4 Mbp; **Supplemental Figure S1**).

Optical maps were generated using the BioNano Genomics technology at 2,057x coverage to further scaffold the Trayshed genome assembly produced previously from Single-Molecule Real-Time (SMRT; Pacific Biosciences) long reads (N50 = 1,479,367 bp; Massonnet *et al.* 2020). The BioNano consensus genome map was combined with the contigs to produce an 896.0 Mbp hybrid assembly nearly twice the expected haploid genome size of Trayshed determined by flow cytometry. Genome sizes estimated by sequencing are known to be underestimates, while the flow cytometry approach tends to produce overestimates (Elliott and Gregory, 2015). In the hybrid assembly, ~ 70% of the complete BUSCO genes were duplicated and, 2.16 ± 1.07 copies of each PN40024 gene were present. Altogether, the size of the assembly, the amount of duplicated BUSCO genes (**Supplemental Table S1**), and the doubled representation of PN20024 genes are strong evidence that the draft assembly represents the diploid genome of Trayshed. Finally, we reconstructed two sets of phased chromosome-scale pseudomolecules (Haplotype 1, Hap1, and Haplotype 2, Hap2; **Table 1**) using a consensus grape genetic map as a guide (Zou *et al.* 2020). The chromosome-scale assemblies were 400.4 Mbp for Hap1 and 369.9 Mbp for Hap2. Based on the flow cytometry data, the whole assembly represents 86.2% of the estimated diploid genome size (2x 483.4 Mbp). On average, chromosome size was 89.09 ± 8.87 % conserved between the two haplotypes (**Supplemental Table S2**). It was not possible to recover the positional information for regions lacking genetic markers, which were therefore excluded from the pseudomolecule assembly (**Table 1**). The unplaced scaffolds represented only 7.5% of the genome assembly, which is much less than in PN40024 V1 (12.4%; Jaillon et al., 2007) and similar to that of the V2 assembly (6%; Canaguier *et al.*, 2017). The unplaced scaffolds were significantly more repetitive than the rest of the genome (52% *vs.* 42%) and carried about 7% of the total protein-coding genes (3,564 genes). The Trayshed assembly also showed a lower level of gaps overall (~ 1 Mbp per haplotype) than PN40024 (~10 Mbp).

**Table 1.** Genome assembly statistics

|  | Haplotype 1 | Haplotype 2 | Unplaced |
| --- | --- | --- | --- |
| Assembly length (bp) | 400,450,509 | 369,985,741 | 62,961,436 |
| Number of scaffolds | 20 | 20 | 1,690 |
| Average length (bp) | 20,022,525 | 18,499,287 | 37,255 |
| Maximum length (bp) | 34,098,201 | 26,289,366 | 1,487,392 |
| N50 length (bp) | 20,338,664 | 18,578,327 | 46,805 |
| Total gap length (bp) | 645,752 | 1,182,032 | 125,041 |
| Repetitive content (%) | 42.54 | 42.35 | 51.99 |
| Number of protein-coding genes | 25,706 | 23,673 | 3,564 |
| Complete BUSCO copies (%) | 90.6 | 86.2 | 9.6 |

The Trayshed assembly structure was confirmed by comparing positions along each chromosome with their linkage group positions within a high-density *M. rotundifolia* genetic map (Lewter *et al*. 2019) (**Supplemental Figures S2**, **S3**, and **S4**). More than 80% of markers’ positions were consistent between the genetic and physical maps, with strong overall correlation (*R* > 0.90) along all chromosomes for both haplotypes.

### Analysis of chromosome structure supports a fusion of chromosomes 7 and 20 in *Vitis*

Each set of Trayshed’s twenty chromosomes was compared with the chromosome-scale Cabernet Sauvignon assembly (Massonnet *et al.*, 2020), which is the most complete and contiguous diploid genome assembly of grapevine currently available. The two genomes were highly syntenic (**Figure 1A**, **Supplemental Figure S5**). Based on the four haplotype comparisons, colinear regions were 2,433 ± 28 bp long and shared 92.19 ± 0.37% identity (**Supplementary Table S3**). On average, short insertions (<50bp) were 40-bp long, long insertions were 1,501-bp long, short deletions were 40-bp, and long deletions were 998-bp. The longest deletion (~50 kbp) was detected on Cabernet Sauvignon chromosome 8 and the longest insertion (~79 kbp) was observed on chromosome 2 (**Supplemental Table S4**).

**Figure 1:**
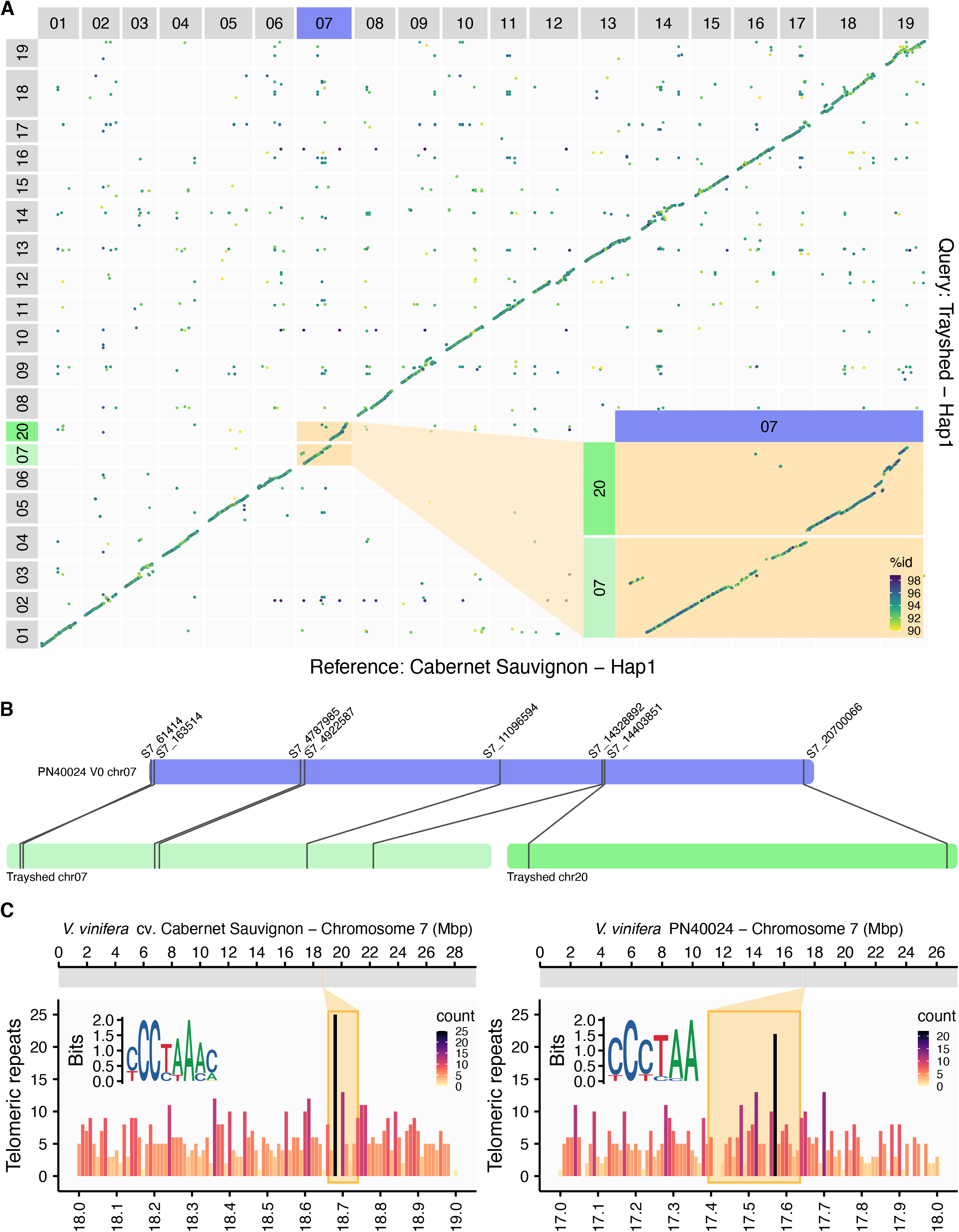
Chromosome fusion localization in *Vitis vinifera*. **A**. Whole-genome alignment of Trayshed Hap1 (y-axis) on Cabernet Sauvignon Hap1 (x-axis). The putative fusion of Trayshed chromosomes 7 and 20 (green) and Cabernet Sauvignon chromosome 7 (purple) is inset. The percentage identity (% id) between the alignments is displayed as a color gradient. **B**. The positions of markers on Trayshed chromosomes 7 and 20 (green) and their corresponding location on PN40024 V0 chromosome 7. **C**. The distribution of telomeric repeats in 1 Mbp containing the expected chromosome fusion in Cabernet Sauvignon (left panel) and PN40024 (right panel). The frequency of telomere repeats is represented as a color gradient. The most enriched motif in the 2-kbp region surrounding the peak of telomeric repeats is inset for each genotype.

Trayshed chromosomes 7 and 20 aligned at the beginning and end of Cabernet Sauvignon chromosome 7, respectively. This supports the fusion of these two chromosomes in *Vitis* reported previously using genetic maps (Blanc *et al.* 2012; Lewter *et al.* 2019) (**Figure 1A**). The marker-containing regions used for Trayshed chromosome structure validation also support this hypothesis. Markers on Trayshed LG 7 and 20 were associated with *V. vinifera* chromosome 7 (**Figure 1B**). Telomeric repeats can be used as indicators of chromosome number reduction rearrangements (Sousa and Renner, 2015). Enrichment of these sequences is expected in a genomic region where a chromosome fusion occurred. To determine the position of chromosome fusion in Cabernet Sauvignon, we searched for telomeric repeats in the region of chromosome 7 that should contain the fusion point based on whole-genome alignments. We examined the genomic region of Cabernet Sauvignon chromosome 7 that spanned Trayshed chromosomes 7 and 20 (chr 7: 18,661,816 - 18,740,440 bp; **Figure 1C**, top panel) and found an enrichment of telomeric repeats in this region (**Figure 1C**, bottom panel). A similar pattern was detected in the hypothetical fusion region of PN40024 (chr 7: 17,392,916 – 17,636,181 bp; **Figure 1C**). Together, these data support a chromosome number reduction by fusion in *V. vinifera* when compared with *M. rotundifolia*.

### Full-length cDNA sequencing and annotation of the protein-coding genes

Together with the public data and RNA-Seq data already available, high-quality full-length cDNA sequences (Iso-Seq; Pacific Biosciences, **Supplemental Figure S6**) were generated as transcriptional evidence to support the gene model predictions. From leaf tissues, 336,932 raw reads were sequenced, clustered, and error corrected with LSC (Au *et al.* 2012) to generate 34,558 high-quality isoforms, 260 low-quality isoforms, and 111,672 singletons (**Supplemental Figure S6**). After obtaining consensus gene models, alternative splice variants were predicted if supported by Iso-Seq data, with 2.87 transcripts per gene on average (**Supplemental Figure S6**). BUSCO genes (Hap1: 92.3%, Hap2: 85.2%) were well-represented in the diploid Trayshed transcriptome. The structural annotation included 52,943 protein-coding gene loci (**Table 1**) and 83,873 proteins (including the alternative forms). There were 25,706 and 23,673 genes on Hap1 and Hap2, respectively (**Table 1**). The repetitive content composed 42.54 and 42.35% of Hap1 and Hap2, respectively, and was higher in the unplaced sequences (51.99%). The coding sequences of each haplotype were aligned on its alternative to evaluate hemizygosity. We found 2,586 and 1,034 hemizygous genes on Hap1 (10.06%) and Hap2 (4.37%), respectively. These results are consistent with *V. vinifera* cv. Zinfandel (4.56%; Vondras *et al.*, 2019) but lower than the levels found in *V. vinifera* cv. Chardonnay (14.6%; Zhou *et al*, 2019).

Based on blastp results (80% identity, 80% coverage), 33,822 (63.88%) Trayshed genes are homologous with 31,471 (54.83%) genes of Cabernet Sauvignon. The 22,755 proteins that did not have a potential ortholog in Cabernet Sauvignon were significantly enriched in functional categories related to cell communication, signal transduction, signaling, regulation of biological and cellular processes, and cellular responses to stimulus and biological regulation (*P* <0.01; Fisher’s exact test). On average, colinear blocks of homologous genes were ~152 genes long (block extension interrupted with more than 25 unmatched genes in a row, **Supplemental Table S5**) based on the synteny results with Cabernet Sauvignon. The longest block of colinear genes was found on chromosome 14 and contained 613 genes (**Supplemental Table S5**).

### NBS-LRR intracellular receptors encoded in the Trayshed genome

Trayshed is attractive to grape breeding programs because it is resistant to pathogens that affect *V. vinifera.* In plants, resistance to pathogens is primarily attributed to the activity of resistance genes (R genes). Nucleotide-binding site leucine-rich repeat (NBS-LRR) genes constitute the largest family of R genes. NBS-LRR genes are associated with resistance to powdery and downy mildew pathogens in several species, including *Arabidopsis* (Warren *et al.* 1998), wheat (He *et al.* 2018), barley (Wei *et al.* 1999; Zhou *et al.* 2001), pepper (Jo *et al.* 2017), and grapes (Riaz *et al.* 2011; Zini *et al.* 2019). We divided the NBS-LRR genes into eight different classes depending on the domains detected in the proteins: CC-NBS-LRR, CC-NBS, CC-X, TIR-NBS-LRR, TIR-NBS, TIR-X, NBS-LRR, NBS (called hereafter NBS-LRR when described as a whole). Overall, the Trayshed genome shows a slightly higher number of NBS-LRR genes (1,158) than Cabernet Sauvignon (1,013). Both species showed the highest number of NBS-LRR localized on chromosomes 9, 12, 13, and 18 (**Figure 2A**, **Supplementary Table S6**). NBS-LRR genes can occur in clusters that could be critical to their function as resistance loci (Hulbert *et al.* 2001; Meyers *et al.* 2003). The average number of clusters detected in Trayshed per haplotype (122) was similar to Cabernet Sauvignon (120), but their arrangement in the genome varied substantially (**Supplemental Figure S7**). Among the haplotypes, in Cabernet Sauvignon, NBS-LRR clusters accumulated the most on chromosome 18 Hap1 while the highest number was observed on chromosome 12 in Trayshed Hap1. These differences could plausibly contribute to disparities in disease resistance.

**Figure 2:**
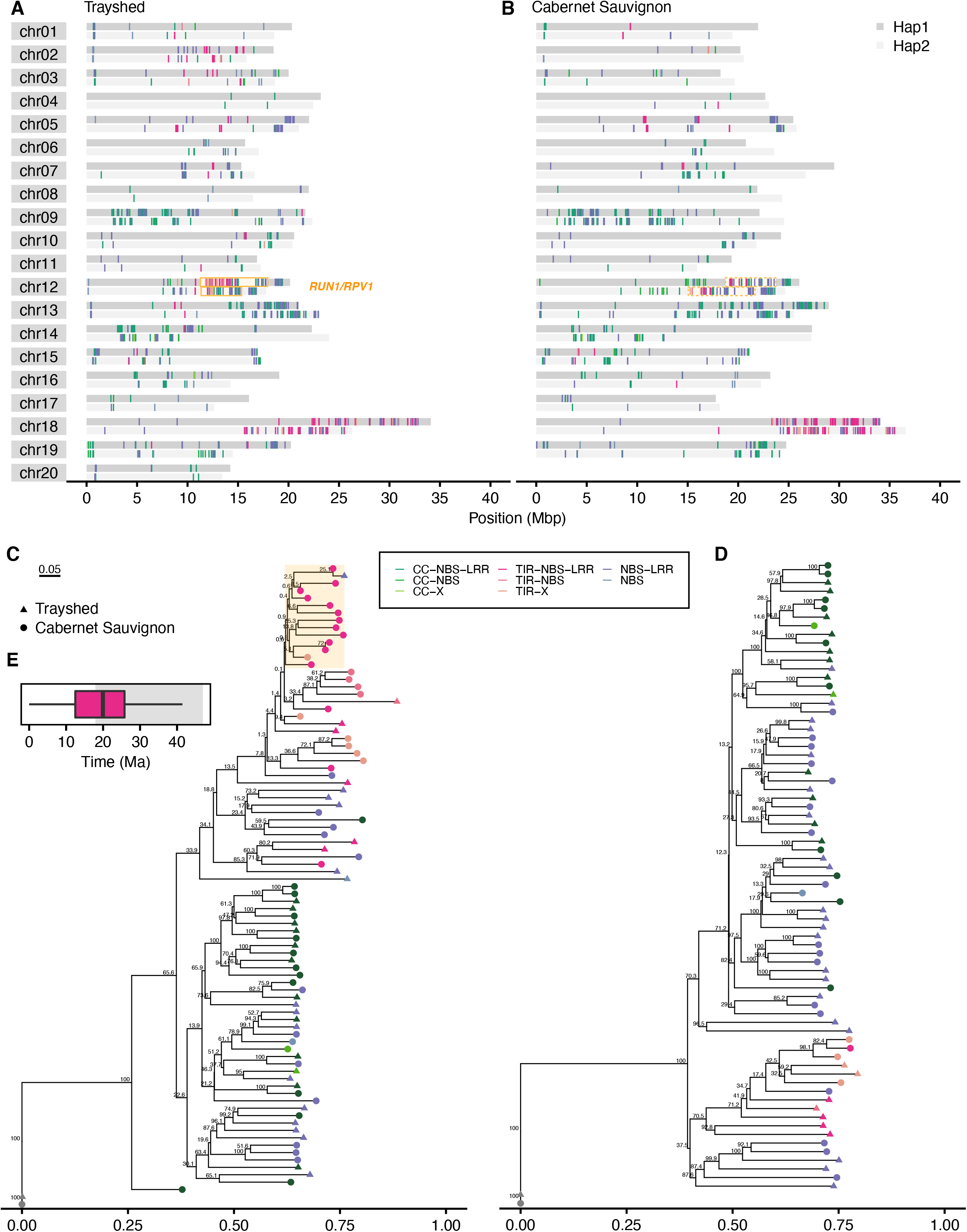
NBS-LRR genes in Trayshed and Cabernet Sauvignon *RUN1/RPV1* loci. Location of the different classes of NBS-LRR along the different chromosomes of the two haplotypes (Hap1: dark grey, Hap2: light grey) in Trayshed (**A**) and Cabernet Sauvignon (**B**). The locations of *RUN1/RPV1* are indicated with an orange box. Phylogenetic trees of NBS-LRR proteins at *RUN1/RPV1* locus in Trayshed and Cabernet Sauvignon, Hap1 (**C**) and Hap2 (**D**). GAPDH was used to root the tree and is colored in grey. **E.** The boxplot represents an approximation of the expansion time of the protein cluster highlighted in orange in panel **C**. The approximation of the divergence time between *Vitis* and *Muscadinia* genera is highlighted in grey. The same color coding is used throughout the figure to distinguish the NBS-LRR classes.

### Expansion of NBS-LRR genes at the *RUN1/RPV1* locus in Trayshed

The clusters of NBS-LRR genes on chromosome 12 in Trayshed are fewer and scattered in Cabernet Sauvignon (**Figures 2A** and **2B**). This chromosome contains a known powdery mildew resistance locus, *RUN1/RPV1* (Pauquet *et al.* 2001), with boundaries defined by the VMC4f3.1 and VMC8g9 SSR markers (Barker *et al.* 2005). This region was studied in Trayshed (Hap1 - chr12: 11,270,914 - 17,943,644 bp; Hap2 – chr12: 11,341,795 - 15,346,610 bp) and Cabernet Sauvignon (Hap1 - chr12: 18,746,148 - 23,727,933 bp; Hap2 – chr12: 15,055,893 - 21,663,411); it included 321 and 201 annotated genes in Trayshed Hap1 and Hap2, respectively, and 287 and 312 genes in Cabernet Sauvignon haplotypes. Within *RUN1/RPV1*, 52 and 33 NBS-LRR were identified in Trayshed Hap1 and Hap2, respectively. For Cabernet Sauvignon, 33 and 40 NBS-LRR were identified in Hap1 and Hap2, respectively.

Notably, most of the NBS-LRR classes are present for both species in this region. A phylogenetic analysis of the NBS-LRR proteins at *RUN1/RPV1* identified a protein cluster specific to Trayshed Hap1 (**Figure 2C**, **Supplemental Figure S8**), absent from the Hap2 (**Figure 2D**). After aligning the coding sequences of those proteins, the synonymous mutation rate was calculated. We estimated that TIR-NBS-LRR genes diverged at nearly 19 Ma (**Figure 2E**). This is one of the latest estimates of the divergence between the *Muscadinia* and *Vitis* genera found in the literature [~18–47 Ma] (Wan *et al.* 2013; Liu *et al.* 2016; Ma *et al.* 2018). Thus, a recent expansion of TIR-NBS-LRR genes might have occurred in only *Muscadinia* and other classes may have expanded prior. To characterize gene duplication in the *RUN1/RPV1* alleles, Cabernet Sauvignon haplotypes were aligned on Trayshed haplotypes to evaluate how many homologous regions are detected using tblastx. Within the resistance locus boundaries of both genomes (and excluding LTR retrotransposons), the highest number of alignments was observed when Trayshed Hap1 was used as the target (**Figure 3**, **Supplemental Figure S9**). There were 10,270 regions aligned with Cabernet Sauvignon Hap1 (**Figure 3A**) and 8,372 alignments between Hap2 (**Figure 3B**). On average, one region in Cabernet Sauvignon Hap1 had 1.75 hits in Trayshed Hap1 and 1.18 hits in Hap2. For the most duplicated region, a single sequence from Cabernet Sauvignon Hap1 matched up to 16 different regions in Trayshed Hap1 (**Figure 3A**). The increased number of hits per window that was observed in Trayshed Hap1 evidenced the expansion of the locus through an increase of the copy number of homologous regions. The regions in Trayshed Hap1 with the highest number of homologs (with at least 10 hits matching a single Cabernet Sauvignon region) were positioned between ~11.83 Mbp and 14.24 Mbp on chromosome 12 (**Figure 3A**) and is similar to the genomic region identified by Feechan *et al.* (2013). It includes UDP-glucosyltransferase-coding genes followed by R gene analogs. In Cabernet Sauvignon, highly duplicated regions were mostly composed of kinetochore NDC80 complex genes; these matched genes with similar or NBS-LRR functions in Trayshed. It is important to note that the assembly of the two *RUN1/RPV1* loci of Trayshed was complex. Despite the currently available long-read sequencing technologies, the highly duplicated content at *RUN1/RPV1* still presents several technical limitations.

**Figure 3:**
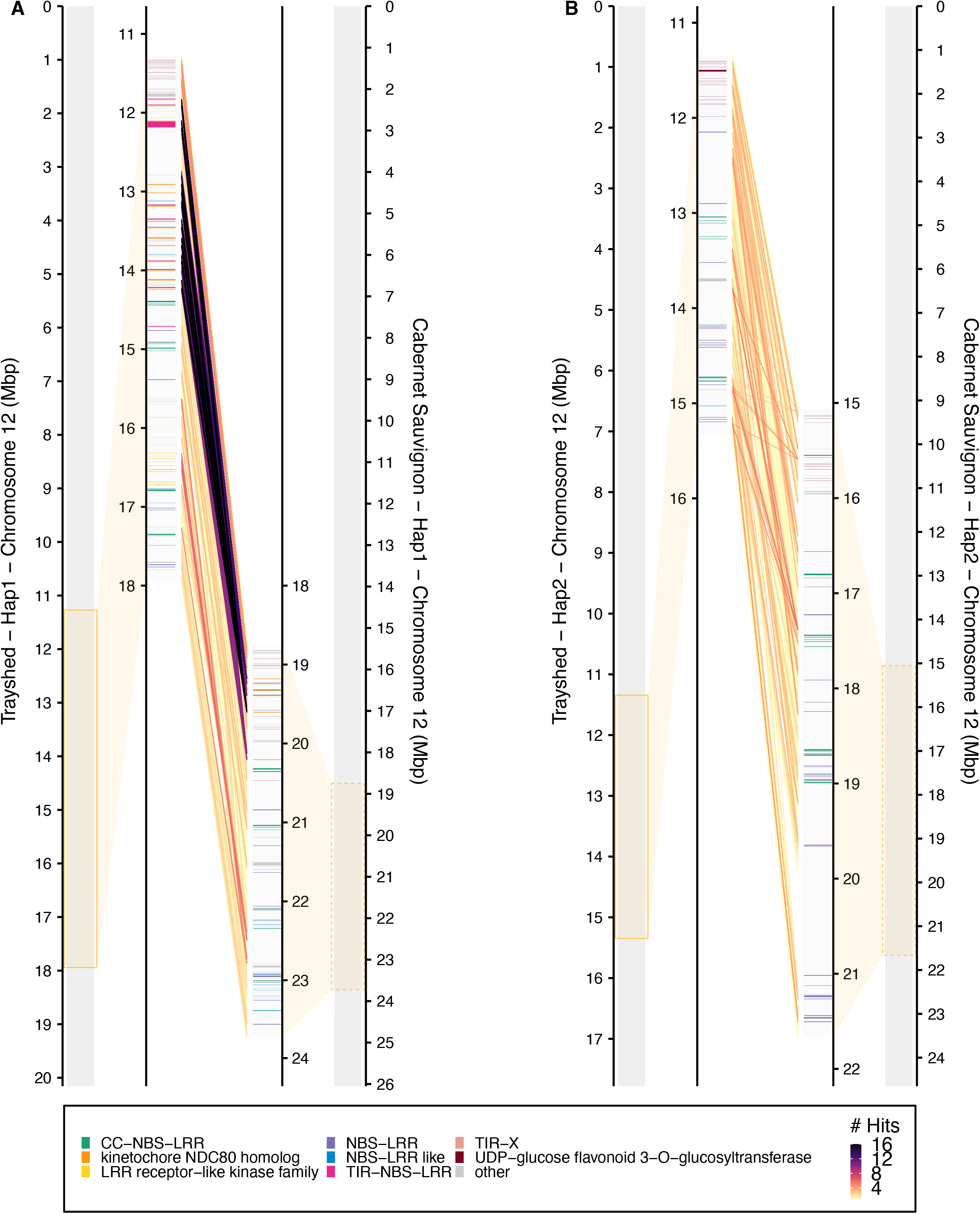
Synteny between the *RUN1/RPV1* alleles in Trayshed and Cabernet Sauvignon. Chromosome 12 of Trayshed (left, target) and Cabernet Sauvignon (right, query) Hap1 (**A**) or Hap2 (**B**) are compared with tblastx of 50Kbp blocks. The *RUN1/RPV1* resistance locus is highlighted with an orange box. Syntenic gene content within the locus is represented as lines in the center of each panel. Duplications of Cabernet Sauvignon sequences are shown using a color gradient. Highly duplicated sequences in Trayshed are dark purple. The functional annotations depicted include categories represented by at least 5 genes. Categories represented by less than 10 genes and not related to disease resistance genes are included in the category, “other”.

Between Trayshed and Cabernet Sauvignon, functionally annotated disease-related genes were differently clustered along the chromosomes. An expansion of TIR-NBS-LRR was identified, which likely occurred after the divergence of their genera. Similar research could be undertaken to characterize the genes or features at other known resistance loci in *M. rotundifolia*, like *RUN2* and *REN5* (Riaz *et al.* 2011; Blanc *et al.* 2012). Acquiring this understanding could be expedited with the availability of high-quality reference sequences for resistant selections, like Trayshed, and susceptible cultivars, like Cabernet Sauvignon and others (Massonnet *et al.*, 2020).

## ACKNOWLEDGMENTS

This work was funded by the NSF grant #1741627 and partially supported by funds to D.C. from the E. & J. Gallo Winery and the Louis P. Martini Endowment in Viticulture.

